# Sequencing and characterization of the complete mitochondrial genome of Critically Endangered Black Soft-shell Turtle (*Nilssonia nigricans*)

**DOI:** 10.1101/361501

**Authors:** Shantanu Kundu, Vikas Kumar, Kaomud Tyagi, Rajasree Chakraborty, Devkant Singha, Iftikar Rahaman, Avas Pakrashi, Kailash Chandra

## Abstract

The complete mitochondrial genome (16796 bp) of an endangered freshwater turtle, *Nilssonia nigricans* was firstly sequenced and annotated. The mitogenome was encoded by 37 genes and a major non-coding Control Region (CR). The mitogenome was A+T biased (62.16%) and spread with six overlapping and 19 intergenic spacer regions. The initiation codons were exceptionally changed as ATA, ATT, and ATC in three Protein-coding genes (PCGs) and a single base (A, T, and C) incomplete termination codons in nine PCGs. The Relative synonymous codon usage (RSCU) value was consistent among all the studied species; exception with significant reduction of Serine (S) frequency in *N. nigricans*, *N. formosa*, and *R. swinhoei*. The secondary structure of *N. nigricans* showed a lack of conventional dihydrouridine (DHU) arm in *trnS* (GCT), as well as formed a small loop structure in the acceptor stem of both *trnR* (TCG) and *trnH* (GTG). The mitogenome of *N. nigricans* also revealed two unique tandem repeats (ATTAT)_8_, and (TATTA)_20_ in CR. Further, the average Ka/Ks values of 13 PCGs were indicating a strong natural selection in the studied Trionychidae species. The constructed Maximum Likelihood (ML) phylogeny by PCGs shows cohesive clustering of *N. nigricans* with *N. formosa*. The resulted phylogeny illustrated the similar topology by all studied species from different taxonomic ranks and supported the previous taxonomic classification. Moreover, further taxon sampling from different taxonomic hierarchy, and their mitogenomics study is vital to reconcile the Testudines phylogeny and assure their evolutionary relationship.

## Introduction

The genus *Nilssonia* (Testudines: Trionychidae) was erected by *N. formosa* as its type and subsequently merged with four *Aspideretes* species^1,2,3^. At present, the five extant *Nilssonia* congeners (*N. formosa*, *N. gangetica*, *N. hurum*, *N. leithii*, and *N. nigricans*) are distributed in India and adjacent countries, from Pakistan, Nepal, Bangladesh, and up to Myanmar. Among them, *N. leithii* is endemic to the southern part of India while *N. formosa* is restricted in Myanmar^4,5^. Further, the identification of *Nilssonia* congeners has been elusive due to the similar conspicuous morphological characters in different life stages^2,3,6,7^. Since the original description, the Black soft-shell turtle, *N. nigricans* was thought to be confined in the shrine pond of Chittagong, Bangladesh^8,9,10^. Due to countless anecdotal reports on its identification and distribution, the species was categorized as ‘Critically Endangered’ up to 2000 by the IUCN SSC Tortoise and Freshwater Turtle Specialist Group (TFTSG). Presently, the status of *N. nigricans* is ‘Extinct in the wild’ according to the International Union for Conservation of Nature (IUCN, Verson-2017-3) Red data list^11^. The species was also included in the Convention on International Trade in Endangered Species of Wild Fauna and Flora (CITES) under ‘Appendix I’ category and recommended to be listed in Indian Wildlife (Protection) Act. In the recent past, the species was categorized as ‘Endangered’ too in the Bangladesh Red List of Reptiles and Amphibians.

As of now, researchers have generated and estimated molecular data to perceive the accurate identification and phylogenetic relationships of *Nilssonia* species^1,4,12,13,14^. The molecular data has also facilitated to resurrect the range distribution of poorly assessed *N. nigricans* in several wild as well as captive localities in northeast India^4,14,15^. Further, the complete mitochondrial genomes add significantly to Protein Coding genes (PCGs), transfer RNA (tRNA), ribosomal RNA (rRNA) and Control Regions (CR), that are useful in systematics and evolutionary research^16,17^. The organization of genes in turtle mitogenome evidenced towards a better understanding of evolution compared with other amniotes (Reptiles, Birds, and Mammals). The genome synteny analysis of turtle mitogenome also depicted the Archosaurian and Diapsid affinity with the sister group of the Archosaurs (Birds and Crocodilians)^18,19^. Furthermore, the gene arrangements in mitogenomes are useful to describe the unusual genomic features and settle phylogenetic hypotheses^20,21,22,23^. So far, the complete mitogenome motifs of *N. nigricans* is anonymous to the world. Thus, in the present study, we sequenced and characterized the complete mitogenome of *N. nigricans*, and compared with other soft-shell turtles (family Trionychidae). Our data would be a valuable resource for further studies on genetic diversity and population structure of *N. nigricans*, which provide a new insights for better conservation strategies in natural settings.

## Materials and methods

### Sample collection and mitochondrial DNA extraction

The survey was conducted with prior permission from the wildlife authority of Arunachal Pradesh Biodiversity Board (Letter No. SFRI/APBB/09-2011-1221-1228 dated 22.07.2016); and all experiments were performed in accordance with relevant guidelines and regulations. The specimen of *N. nigricans* was collected from the tributaries of Brahmaputra River in Arunachal Pradesh state (Longitude 27°45’ N and latitude 95°48’ E) in northeast India (Fig. S1). The blood sample was collected aseptically by micro-syringe from the hind limb and stored in EDTA vial at 4°C. After collecting the blood sample as reliable source of DNA, the specimen was released back in the same eco-system with sufficient care. To remove nuclei and cell debris, a drop of blood sample was centrifuged at 700X g for 5 min at 4°C in 1 ml buffer (0.32 M Sucrose, 1 mM EDTA, 10 mM TrisHCl). The supernatant was collected in 1.5 ml centrifuge tubes and centrifuged at 12000X g for 10 min at 4°C to pellet intact mitochondria. The mitochondrial pellet was suspended in 200 μl of lysis buffer (50 mM TrisHCl, 25 mM of EDTA, 150 mM NaCl), with the addition of 10 μl of proteinase K (20 mg/ml) followed by incubation at 56 °C for 1 hour. Finally, the mitochondrial DNA was purified by Qiagen DNeasy Blood & Tissue Kit. The extracted DNA quality was assessed by electrophoresis in a 1% agarose gel stained with ethidium bromide and final concentration was quantified by NANODROP 2000 spectrophotometer (Thermo Scientific, USA). The mitochondrial DNA was deposited in Centre for DNA Taxonomy laboratory, Zoological Survey of India, Kolkata under voucher IDs ‘ZSI_NFGR-TT8’.

### Genome sequencing, assembly and annotation

The complete mitochondrial genome sequencing, assembly and annotation were carried out at the Genotypic Technology Pvt. Ltd. Bangalore, India (http://www.genotypic.co.in/). For library assembly, 200 ng of DNA was used in the Illumina TruSeq Nano DNA HT library preparation kit. After the fragmentation of mitochondrial DNA by ultra-sonication (Covaris M220, Covaris Inc., Woburn, MA, USA), the purified A-tailed fragments were ligated with the sequencing indexed adapters. The fragments with an insert size of 450 bp were selected using sample purification beads and amplified by Polymerase chain reaction (PCR) to enrich it. The amplified PCR library was analyzed by Bioanalyzer 2100 (Agilent Technologies, Inc., Waldbronn, Germany) using High Sensitivity DNA chips. After obtaining the required concentration and mean peak size, the library was subjected to sequencing in Illumina HiSeq 2000 using 2× 150 bp kit (Illumina, Inc, USA). The high quality paired ends reads were further assembled to generate the contigs in Geneious v.6.0.6 (www.geneious.com) using default parameters^24^. The .fasta-formatted mitochondrial genome assembly was performed by aligning the contigs against the non-redundant nucleotide database of GenBank using the BLASTn search algorithm (http://blast.ncbi.nlm.nih.gov/Blast). The sequence annotation was also checked in MITOS online server (http://mitos.bioinf.uni-leipzig.de). The nucleotide sequences of the PCGs were initially translated into the putative amino acid sequences on the basis of the vertebrate mitochondrial DNA genetic code. The exact initiation and termination codons were identified in ClustalX using other reference sequences of Testudines^25^.

### Genome sequence and Phylogenetic analysis

The direction of locus, size, start and stop codon of PCGs, overlapping regions and intergenic spacers between genes, anticodon of tRNA genes were checked in MITOS online server and Open Reading Frame Finder (https://www.ncbi.nlm.nih.gov/orffinder/). The PCGs situated in light strand (L strand) were made reverse complementary before incorporating them in the analysis. However, the nucleotide composition and ATGC- Skew of tRNA genes, situated in both heavy strand (H strand) and light strand (L strand) were analyzed as its original orientation. The CGView Server (http://stothard.afns.ualberta.ca/cgview_server/) with default parameters was used to map the circular representation of the generated completed mitochondrial genome^26^. The base composition of nucleotide sequences, and composition skewness were calculated manually in Microsoft Excel as described previously: AT skew = [A–T]/[A+T], GC skew = [G–C]/[G+C]^27^. The Relative Synonymous Codon Usage (RSCU) values of each PCGs were calculated by assuring their codon frame using MEGA6.0^28^. The tRNA genes were verified in MITOS online server (http://mitos.bioinf.uni-leipzig.de), tRNAscan-SE Search Server 2.0 (http://lowelab.ucsc.edu/tRNAscan-SE/) and ARWEN 1.2 with the default settings with the appropriate anticodon capable of folding into the typical cloverleaf secondary structure^29,30^. The large and small subunit of RNA (rrnL and rrnS) were annotated by the MITOS online server (http://mitos.bioinf.uni-leipzig.de). The tandem repeats and unique base composition in the control regions were predicted by the online Tandem Repeats Finder web tool (https://tandem.bu.edu/trf/trf.html)^31^. The software package DnaSPv5.0 was used to calculate the synonymous substitutions per synonymous sites (Ks) and non-synonymous substitutions per non-synonymous sites (Ka)^32^. The twelve Trionychidae turtles mitogenomes were acquired from GenBank and incorporated in the dataset for comparative analysis. Further, the complete mitogenomes of five turtle species under five different families (suborder: Cryptodira) were also incorporated to the dataset to illustrate the phylogenetic relationships. The complete mitogenome of *Chelus fimbriata*, family Chelidae (suborder: Pleurodira) was incorporated as an out-group in the phylogenetic analysis. Each PCGs was aligned individually by codons using MAFFT algorithm implemented in TranslatorX with L-INS-i strategy and default settings^33^. To concatenated the PCGs and removed the poorly unaligned sites, GBlocks v0.91 in TranslatorX (http://translatorx.co.uk/) was used with default settings^34^. To construct the Maximum Likelihood (ML) tree, the best model ‘GTR+G+I’ was selected with the lowest Bayesian information criterion (BIC) scores (158167.27) by using Maximum Likelihood statistical method in MEGA6.0 program^28^.

## Results and discussion

### Genome size and organization

In this study, the complete mitogenomes (16796 bp) of the black soft-shell turtle, *N. nigricans* was firstly determined under the ‘National Faunal Genome Resources (NFGR)’ program at the Zoological Survey of India, Kolkata (Fig. 1). The generated mitogenome was submitted in the GenBank database through Sequin submission tool (https://www.ncbi.nlm.nih.gov/proiects/Sequin/) and acquired the accession number: MG383833. The mitogenome encodes for 37 genes; 13 proteins, 22 tRNAs, two rRNAs, and a major non-coding control region (Table 1). Among them, 28 genes (12 PCGs, 2 rRNA genes and 14 tRNA genes) were located on the heavy strand (H strand) and others (*nad6* and eight tRNA genes) were located on the light strand (L strand). The gene rearrangement of *N. nigricans* is same as described for the typical vertebrate species^35^. The gene arrangements in both L and H strand were also similar in other Trionychidae species (Table S1).

**Figure 1.**
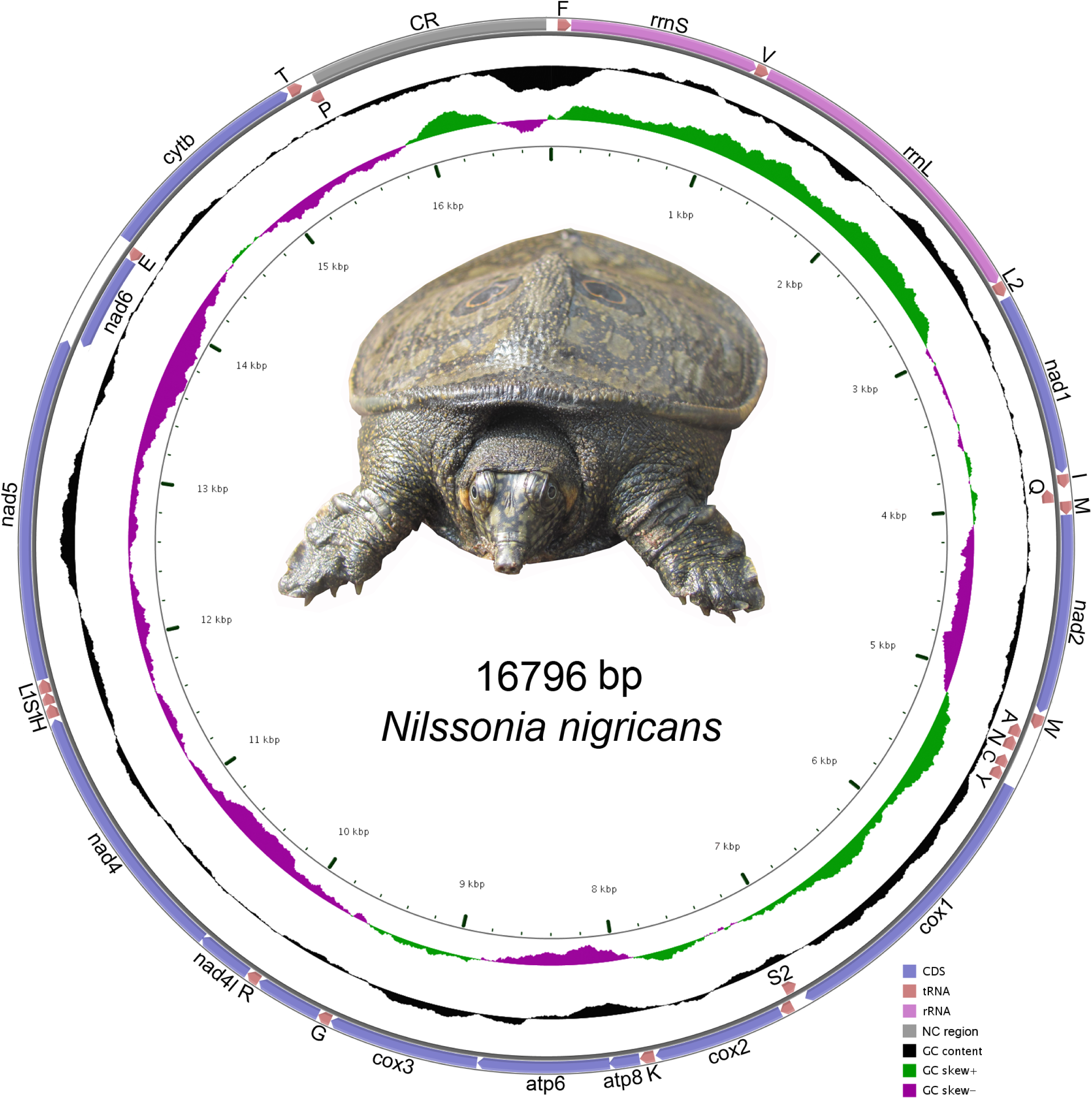
The mitochondrial genome of *N. nigricans*. Direction of gene transcription is indicated by arrows. PCGs are shown as blue arrows, rRNA genes as orchid arrows, tRNA genes as coral arrows and 1290 bp non coding region as grey rectangle. tRNAs are encoded according to their single-letter abbreviations. The GC content is plotted using a black sliding window, GC-skew is plotted using green and violet color sliding window as the deviation from the average in the complete mitogenome. The figure was drawn using CGView online server (http://stothard.afns.ualberta.ca/cgview_server/) with default parameters. The Species photographs was taken by the first authors (S.K.) by using Nikon D3100 and edited manually in Adobe Photoshop CS 8.0.

**Table 1.** List of annotated mitochondrial genes of *Nilssonia nigricans*.

| Locus Name | Direction | Location | Size (bp) | Anti codon | Start codon | Stop codon | Intergenic Nucleotides |
| --- | --- | --- | --- | --- | --- | --- | --- |
| *trnF* | + | 59-128 | 70 | GAA | - | - | 0 |
| *rrnS* | + | 129-1110 | 982 | - | - | - | -2 |
| *trnV* | + | 1109-1178 | 70 | TAC | - | - | 0 |
| *rrnL* | + | 1179-2789 | 1611 | - | - | - | 0 |
| *trnL2* | + | 2790-2865 | 76 | TAA | - | - | 13 |
| *nad1* | + | 2879-3823 | 945 | - | ATA | A | 8 |
| *trnI* | + | 3832-3901 | 70 | GAT | - | - | -1 |
| *trnQ* | - | 3901-3971 | 71 | TTG | - | - | -1 |
| *trnM* | + | 3971-4039 | 69 | CAT | - | - | 0 |
| *nad2* | + | 4040-5068 | 1029 | - | ATG | A | 13 |
| *trnW* | + | 5082-5156 | 75 | TCA | - | - | 4 |
| *trnA* | - | 5161-5229 | 69 | TGC | - | - | 1 |
| *trnN* | - | 5231-5304 | 74 | GTT | - | - | 33 |
| *trnC* | - | 5338-5403 | 66 | GCA | - | - | 0 |
| *trnY* | - | 5404-5471 | 68 | GTA | - | - | 4 |
| *cox1* | + | 5476-7008 | 1533 | - | ATT | A | 4 |
| *trnS2* | - | 7013-7083 | 71 | TGA | - | - | 0 |
| *trnD* | + | 7084-7152 | 69 | GTC | - | - | 0 |
| *cox2* | + | 7153-7833 | 681 | - | ATG | T | 7 |
| *trnK* | + | 7841-7916 | 76 | TTT | - | - | 1 |
| *atp8* | + | 7918-8076 | 159 | - | ATG | A | -4 |
| *atp6* | + | 8073-8753 | 681 | - | ATG | (TAA) | 2 |
| *cox3* | + | 8756-9538 | 783 | - | ATG | T | 1 |
| *trnG* | + | 9540-9609 | 70 | TCC | - | - | 0 |
| *nad3* | + | 9610-9958 | 348 | - | ATG | AGA | 1 |
| *trnR* | + | 9960-10030 | 71 | TCG | - | - | 1 |
| *nad4l* | + | 10032-10325 | 294 | - | ATG | (TAA) | -4 |
| *nad4* | + | 10322-11692 | 1371 | - | ATG | (C) | 10 |
| *trnH* | + | 11703-11772 | 70 | GTG | - | - | 0 |
| *trnS1* | + | 11773-11834 | 62 | GCT | - | - | -1 |
| *trnL1* | + | 11834-11905 | 72 | TAG | - | - | 0 |
| *nad5* | + | 11906-13681 | 1776 | - | ATG | (TAA) | 1 |
| *nad6* | - | 13683-14204 | 522 | - | ATG | (T) | 0 |
| *trnE* | - | 14205-14272 | 68 | TTC | - | - | 9 |
| *cytb* | + | 14282-15409 | 1128 | - | ATC | A | 9 |
| *trnT* | + | 15419-15492 | 74 | TGT | - | - | 2 |
| *trnP* | - | 15495-15564 | 70 | TGG | - | - | 0 |
| A+T-rich Region |  | 15565-16796<br>1-58 | 1290 | - | - | - | - |

The total length of PCGs, tRNAs, and rRNAs was 11251 bp, 1551 bp, and 2593 bp respectively. The nucleotide composition was biased toward A+T (62.16%) within the complete mitogenome of *N. nigricans*. The A+T composition was 61.26%, 63.82%, 61.47%, and 68.52% in PCGs, tRNAs, rRNAs and CR respectively (Table 2). In other Trionychidae species, the nucleotide composition within the complete mitogenomes is similar to *N. nigricans* and biased towards A+T with a variable frequency ranging from 58.48% (*T. triunguis*) to 62.97% (*N. formosa*). The AT skewness was 0.197 and GC skewness was −0.400, resulted in the complete mitogenome of *N. nigricans*. The AT skewness of other Trionychidae species were varied from 0.127 (*P. sinensis*) to 0.208 (*T. triunguis*) and GC skewness were varied from −0.412 (*T. triunguis*) to −0.363 (*P. steindachneri*). The resulted positive AT skew in most of the genes indicated more Adenine (A)s than Thymine (T)s in the complete genome of Trionychidae species (Table 2).

**Table 2.** Nucleotide composition of the mitochondrial genome in different Trionychid turtle's mtDNA. The A+T biases of whole mitogenome, protein coding genes, tRNA, rRNA, and control regions were calculated by AT-skew = (A-T)/(A+T) and GC-skew= (G-C)/(G+C), respectively.

| Species | Size | A% | T% | G% | C% | A+T% | AT- Skew | GC- Skew |
| --- | --- | --- | --- | --- | --- | --- | --- | --- |
| **Complete mitogenome** |  |  |  |  |  |  |  |  |
| *A. ferox* | 16866 | 35.80 | 25.38 | 11.82 | 26.98 | 61.18 | 0.170 | -0.390 |
| *N. nigricans* | 16796 | 37.20 | 25.00 | 11.30 | 26.50 | 62.20 | 0.196 | -0.402 |
| *A. spinifera* | 16749 | 35.52 | 25.26 | 12.09 | 27.12 | 60.78 | 0.168 | -0.383 |
| *C. indica* | 16726 | 35.79 | 24.33 | 11.87 | 27.99 | 60.12 | 0.190 | -0.404 |
| *D. subplana* | 17289 | 35.29 | 25.96 | 11.94 | 26.78 | 61.26 | 0.152 | -0.383 |
| *L. punctata* | 16489 | 35.73 | 25.68 | 12.22 | 26.36 | 61.41 | 0.163 | -0.366 |
| *L. scutata* | 16512 | 35.97 | 25.41 | 12.04 | 26.56 | 61.38 | 0.172 | -0.376 |
| *N. formosa* | 17145 | 37.13 | 25.83 | 11.19 | 25.82 | 62.97 | 0.179 | -0.395 |
| *P. steindachneri* | 17243 | 35.24 | 26.51 | 12.28 | 26.34 | 61.36 | 0.135 | -0.363 |
| *P. cantorii* | 17499 | 35.24 | 24.21 | 11.98 | 28.54 | 59.46 | 0.185 | -0.408 |
| *P. sinensis* | 17364 | 35.22 | 27.26 | 11.77 | 25.73 | 62.49 | 0.127 | -0.372 |
| *R. swinhoei* | 16990 | 34.98 | 25.46 | 12.24 | 27.30 | 60.45 | 0.157 | -0.380 |
| *T. triunguis* | 16590 | 35.338 | 23.14 | 12.20 | 29.31 | 58.48 | 0.208 | -0.412 |
| **Protein Coding genes (PCG)** |  |  |  |  |  |  |  |  |
| *A. ferox* | 11359 | 35.32 | 24.91 | 10.68 | 29.07 | 60.23 | 0.172 | -0.462 |
| *N. nigricans* | 11251 | 36.60 | 28.60 | 10.13 | 24.66 | 65.20 | 0.122 | -0.417 |
| *A. spinifera* | 11350 | 34.70 | 24.88 | 11.09 | 29.32 | 59.58 | 0.164 | -0.451 |
| *C. indica* | 11360 | 35.38 | 24.02 | 10.62 | 29.96 | 59.41 | 0.191 | -0.476 |
| *D. subplana* | 11335 | 35.13 | 25.77 | 10.82 | 28.26 | 60.90 | 0.153 | -0.446 |
| *L. punctata* | 11342 | 35.18 | 25.98 | 10.80 | 28.02 | 61.17 | 0.150 | -0.443 |
| *L. scutata* | 11356 | 35.37 | 25.73 | 10.68 | 28.20 | 61.11 | 0.157 | -0.450 |
| *N. formosa* | 11357 | 36.98 | 25.12 | 10.05 | 27.84 | 62.10 | 0.190 | -0.469 |
| *P. steindachneri* | 11216 | 34.65 | 26.25 | 10.81 | 28.27 | 60.91 | 0.137 | -0.446 |
| *P. cantorii* | 11357 | 34.49 | 23.36 | 11.09 | 31.04 | 57.85 | 0.192 | -0.473 |
| *P. sinensis* | 11385 | 35.26 | 27.69 | 10.52 | 26.51 | 62.96 | 0.120 | -0.431 |
| *R. swinhoei* | 11384 | 34.59 | 25.14 | 11.09 | 29.16 | 59.74 | 0.158 | -0.448 |
| *T. triunguis* | 11352 | 34.39 | 22.87 | 11.17 | 31.55 | 57.26 | 0.201 | -0.476 |
| **tRNA genes** |  |  |  |  |  |  |  |  |
| *A. ferox* | 1602 | 35.95 | 27.65 | 14.29 | 22.09 | 63.6 | 0.13 | -0.214 |
| *N. nigricans* | 1551 | 36.49 | 27.33 | 14.18 | 21.98 | 63.82 | 0.14 | -0.22 |
| *A. spinifera* | 1594 | 35.57 | 27.41 | 14.74 | 22.27 | 62.98 | 0.129 | -0.203 |
| *C. indica* | 1601 | 35.47 | 26.92 | 14.61 | 22.98 | 62.39 | 0.137 | -0.222 |
| *D. subplana* | 1586 | 35.81 | 27.23 | 14.5 | 22.44 | 63.05 | 0.136 | -0.215 |
| *L. punctata* | 1600 | 35.37 | 26.68 | 14.87 | 23.06 | 62.06 | 0.139 | -0.215 |
| *L. scutata* | 1596 | 34.89 | 26.56 | 15.22 | 23.3 | 61.46 | 0.135 | -0.209 |
| *N. formosa* | 1600 | 36.5 | 27.62 | 14.31 | 21.56 | 64.12 | 0.138 | -0.202 |
| *P. steindachneri* | 1589 | 35.3 | 27.69 | 15.1 | 21.9 | 62.99 | 0.12 | -0.183 |
| *P. cantorii* | 1591 | 34.63 | 25.89 | 15.27 | 24.19 | 60.52 | 0.144 | -0.226 |
| *P. sinensis* | 1617 | 35.37 | 28.57 | 14.34 | 21.7 | 63.94 | 0.106 | -0.204 |
| *R. swinhoei* | 1586 | 35.18 | 27.23 | 14.81 | 22.76 | 62.42 | 0.127 | -0.211 |
| *T. triunguis* | 1587 | 35.47 | 26.08 | 14.87 | 23.56 | 61.56 | 0.152 | -0.226 |
| **rRNA genes** |  |  |  |  |  |  |  |  |
| *A. ferox* | 2584 | 40.05 | 21.20 | 15.86 | 22.87 | 61.26 | 0.307 | -0.180 |
| *N. nigricans* | 2593 | 40.18 | 21.28 | 15.77 | 22.75 | 61.47 | 0.307 | -0.181 |
| *A. spinifera* | 2585 | 40.11 | 21.04 | 16.05 | 22.78 | 61.16 | 0.311 | -0.173 |
| *C. indica* | 2570 | 39.37 | 20.38 | 16.53 | 23.69 | 59.76 | 0.317 | -0.177 |
| *D. subplana* | 2578 | 39.13 | 21.72 | 16.13 | 23.00 | 60.86 | 0.286 | -0.175 |
| *L. punctata* | 2547 | 38.43 | 21.94 | 16.92 | 22.69 | 60.38 | 0.273 | -0.145 |
| *L. scutata* | 2585 | 39.38 | 21.47 | 16.28 | 22.86 | 60.85 | 0.294 | -0.167 |
| *N. formosa* | 2594 | 40.28 | 21.81 | 15.57 | 22.32 | 62.10 | 0.297 | -0.178 |
| *P. steindachneri* | 2585 | 39.30 | 21.27 | 16.20 | 23.21 | 60.58 | 0.297 | -0.177 |
| *P. cantorii* | 2567 | 39.65 | 20.17 | 16.40 | 23.76 | 59.83 | 0.325 | -0.183 |
| *P. sinensis* | 2687 | 39.18 | 22.55 | 15.96 | 22.29 | 61.74 | 0.269 | -0.165 |
| *R. swinhoei* | 2573 | 38.63 | 21.33 | 16.75 | 23.28 | 59.96 | 0.288 | -0.163 |
| *T. triunguis* | 2586 | 39.24 | 19.76 | 16.04 | 24.94 | 59.01 | 0.330 | -0.216 |
| **Control regions (CR)** |  |  |  |  |  |  |  |  |
| *A. ferox* | 1357 | 31.61 | 35.74 | 10.31 | 22.32 | 67.35 | -0.061 | -0.367 |
| *N. nigricans* | 1290 | 36.66 | 31.86 | 9.84 | 21.62 | 68.52 | 0.07 | -0.374 |
| *A. spinifera* | 1234 | 33.22 | 36.14 | 9.23 | 21.39 | 69.36 | -0.042 | -0.396 |
| *C. indica* | 1227 | 33 | 32.92 | 10.1 | 23.96 | 65.93 | 0.001 | -0.406 |
| *D. subplana* | 1820 | 30.54 | 32.91 | 10.6 | 25.93 | 63.46 | -0.037 | -0.419 |
| *L. punctata* | 1008 | 34.92 | 31.74 | 11.8 | 21.52 | 66.66 | 0.047 | -0.291 |
| *L. scutata* | 1005 | 35.22 | 31.24 | 11.64 | 21.89 | 66.46 | 0.059 | -0.305 |
| *N. formosa* | 1604 | 33.97 | 36.34 | 8.54 | 21.13 | 70.32 | -0.033 | -0.424 |
| *P. steindachneri* | 1736 | 29.37 | 36.34 | 13.19 | 21.08 | 65.72 | -0.106 | -0.23 |
| *P. cantorii* | 1994 | 34.65 | 33.75 | 8.62 | 22.96 | 68.4 | 0.013 | -0.453 |
| *P. sinensis* | 1830 | 29.5 | 31.03 | 10.87 | 28.57 | 60.54 | -0.025 | -0.448 |
| *R. swinhoei* | 1471 | 31.67 | 34.46 | 10.12 | 23.72 | 66.14 | -0.042 | -0.401 |
| *T. triunguis* | 1078 | 35.8 | 31.26 | 9.18 | 23.74 | 67.06 | 0.067 | -0.442 |

### Overlapping and intergenic spacer regions

Six overlapping sequences with a total length of 13 bp were identified in the *N. nigricans* complete mitogenome (Table 1). These sequences varied in length from 1 bp to 4 bp with the longest overlapping region present between NADH dehydrogenase subunit 4L (*nad4l*) and NADH dehydrogenase subunit 4 (*nad4*) as well as in between ATP synthase F0 subunit 8 (*atp8*) and ATP synthase F0 subunit 6 (*atp6*). The intergenic spacers within this mitogenomes, spread over 19 regions and ranged in size from 1 bp to 33 bp with a total length of 124 bp. The longest spacer (33 bp) was occurred between *trnN* (GTT) and *trnC* (GCA). In other Trionychidae species, the number of overlapping sequences were varied from five (*N. formosa*) to 14 (*P. sinensis*) with a length variation of 7 bp to 143 bp, respectively with longest overlapping region present between the small subunit of ribosomal RNA (*rrnS*) and *trnV* (TAC). The longest intergenic spacer (33bp) is present between *trnN* (GTT) and *trnC* (GCA) of *L. scutata* and *L. punctata* (Table S2).

### Protein-coding gene (PCG) features

The PCGs region of *N. nigricans* was 11251 bp long and shares 66.9% of the complete mitogenome. The contrast of nucleotide composition, AT skew and GC skew of PCGs of all Trionychidae species were exhibited in Table 2. The AT skew value of the PCGs was 0.136 and the GC skew value of the PCGs was −0.414 in *N. nigricans*. The AT skew value of other Trionychidae species PCGs varied from 0.069 (*P. sinensis*) to 0.138 (*T. triunguis*) and GC skew value from −0.450 (*L. scutata*) to −0.370 (*P. sinensis*) (Table 2). Further, the PCGs of *N. nigricans* start with ATG initiation codon, except for NADH dehydrogenase subunit 1 (*nad1*) with ATA, Cytochrome oxidase subunit I (*cox1*) with ATT, and Cytochrome b (*cytb*) with ATC.

In *N. nigricans*, out of 13 PCGs, three PCGs (*atp6*, *nad4l*, *nad5*) used a typical (TAA) termination codon; whereas the remaining three PCGs (*cox2*, *cox3*, *nad6*) terminated with a single base (T), five PCGs (*nad1*, *nad2*, *cox1*, *atp8*, *cytb*) terminated with a single base (A), *nad3* terminated with AGA, and *nad4* terminated with a single base (C) (Table 1). In other Trionychidae species, most of the start codon for PCGs was ATG, except for ATT (*cox1* of *N. formosa*), ATC (*cytb* of *N. formosa*), GTG (*cox1* of *D. subplana*, *P. steindachneri*, *P. sinensis*, *R. swinhoei*, *A. spinifera*, *A. ferox*, *T. triunguis*, *C. indica*, *P. cantorii*, *L. scutata*, *L. punctata*; and *nad5* of *L. scutata*), ATA (*nad1* and *nad6* of *P. steindachneri*; *nad6* of *A. spinifera*, *A. ferox*, *C. indica*), ATT (*cytb* of *P. sinensis*). The termination codon was TAA for most of the Trionychidae species, except for single base (T): *nad1* of *N. formosa; cox3* and *nad3* of *N. formosa*, *D. subplana*, *P*. steindachneri, *A. spinifera*, *R. swinhoei*, *A. ferox*, *C. indica*, *P. cantorii*, *L. scutata*, *L. punctata*; *nad2* and *nad4* of *P. steindachneri*, *A. spinifera*, *A. ferox*, *T. triunguis*, *C. indica*, *P. cantorii*, *L. scutata*, *L. punctata; nad4* of *D. subplana*, *R. swinhoei; nad4l* of *D. subplana; cytb* of *R. swinhoei*, *A. ferox; cox3* of *T. triunguis*. The termination codon (TA) was observed in *nad1* of *P. steindachneri*, *R. swinhoei*, *A. spinifera*, *A. ferox; nad2* and *nad4l* of *R. swinhoei*. The termination codon AGA was observed in *nad6* of *D. subplana*, *C. indica*, *L. punctata*, *P. steindachneri*, *R. swinhoei*, *A. spinifera*, *A. ferox*, *P. cantorii*, *L. scutata*, *T. triunguis*; *cox1* of *P. steindachneri*, *R. swinhoei*, *A. spinifera*, *A. ferox*, *P. cantorii*, *L. scutata*, *P. sinensis; nad3* of *T. triunguis*. The termination codon AGG was observed in *cox1* of *C. indica; nad6* of *P. sinensis*. The termination codon TAG was observed in *nad1* of *D. subplana*, *P. sinensis*, *L. punctata; nad2* of *D. subplana*, *P. sinensis; nad5* of *L. scutata* (Table S3). The codons of each amino acids were conserved in all PCGs of the compared Trionychidae species. Moreover, codons with A or T in the third position were overused in comparison to other synonymous codons. For example, the codon for Glutamine (Gln) as CAG was rare as compared to CAA; and codon for Glutamic acid (Glu) as GAG was rare as compared to GAA. The RSCU analysis indicated a significant fall in Serine (Ser) frequency in *N. nigricans*, *N. formosa*, and *R. swinhoei* (Table S4, Fig. 2). Relative synonymous codon usage (RSCU) analysis of PCGs in N. *nigricans* revealed that the codons encoding Asn, His, Ile, Leu, Met, Thr, Pro and Ser were the most frequently present, nevertheless those encoding Arg, Asp, Cys and Glu were rare (Fig. 3). RSCU analysis of other Trionychidae species showed the same frequency of codon usage for their respective amino acids.

**Figure 2.**
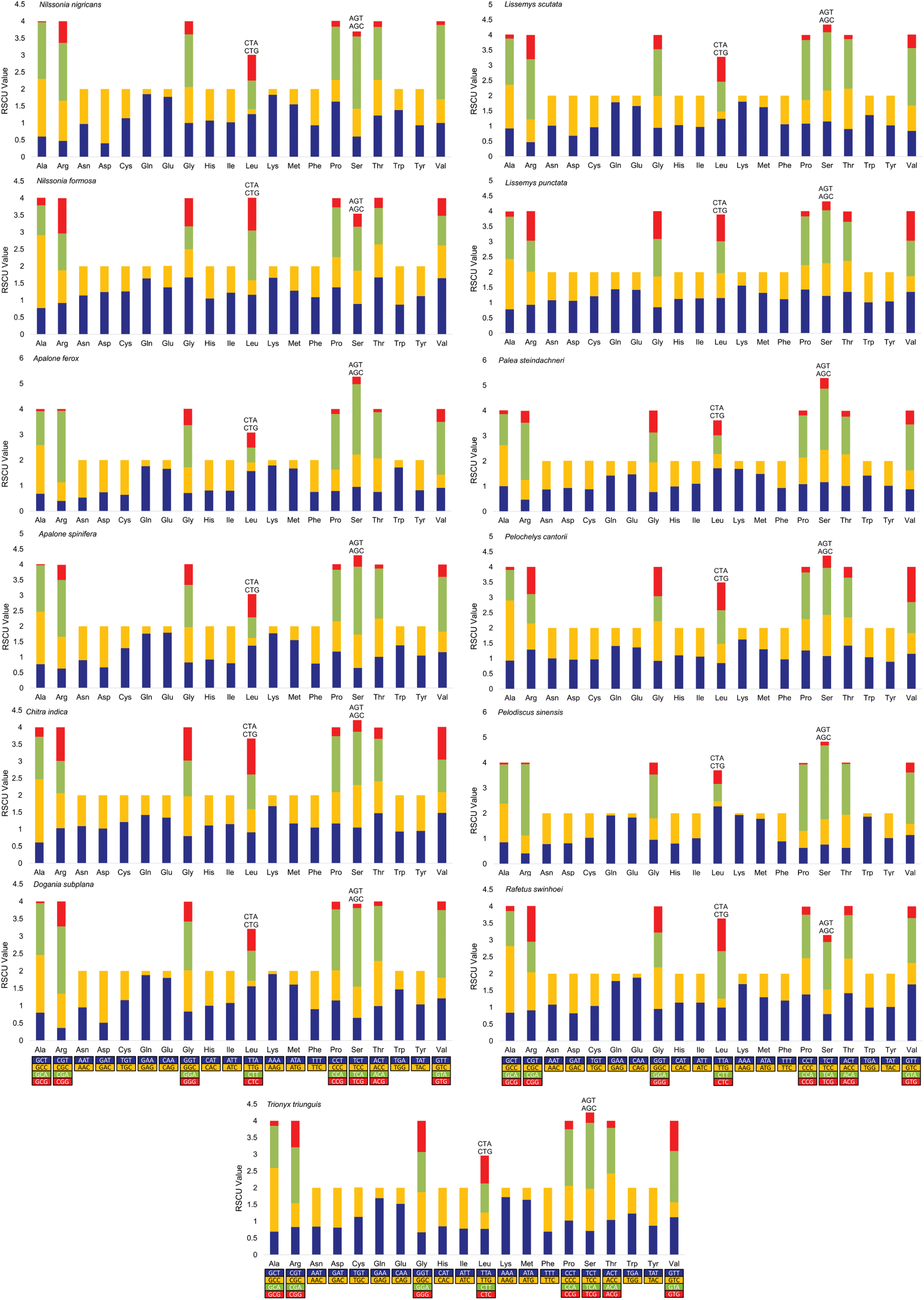
Relative synonymous codon usage (RSCU) in the mitogenome of *N. nigricans* and comparison with other Trionychidae species. The cumulative RSCU values were represented on the y-axis while the codon families for each amino acid are represented on the x-axis. Codons more than four combination are specified at the top of the corresponding amino acids columns. Amino acids are encoded according to their three-letter abbreviations. The figure was drawn using MEGA6, and edited manually in Microsoft Excel.

**Figure 3.**
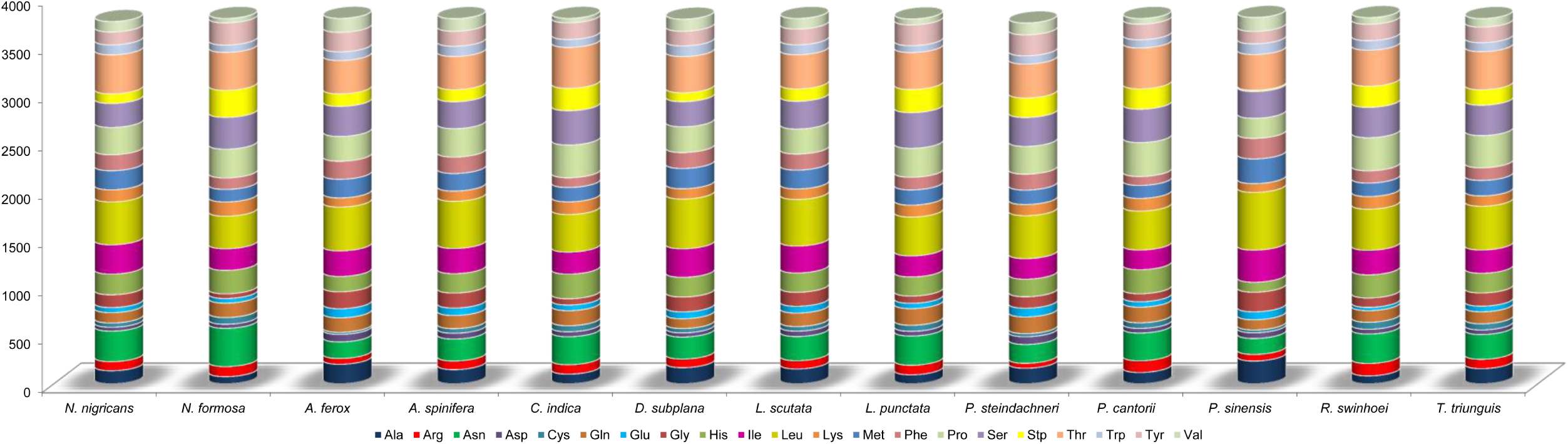
Comparison of codon usage within the mitochondrial genome of members of the Trionychidae species.

### Transfer RNAs and ribosomal RNAs

The representative forecasted structures of 22 tRNAs total of 1551 bp were identified for *N. nigricans* complete mitogenome, ranging from 62 bp to 76 bp. Among them, 14 were present in major strand and remaining eight were present in minor strand (Table 1). In other Trionychidae species, the total length of tRNA genes was varied from 1540 bp (*D. subplana* and *R. swinhoei*) to 1571 bp (*P. sinensis*). The AT skewness of tRNAs in *N. nigricans* was 0.143 and GC skewness was −0.215. In other species, AT skewness was varied from 0.111 (*P. sinensis*) to 0.159 (*T. triunguis*) and GC skewness was varied from −0.226 (*P. cantorii and T. triunguis*) to −0.184 (*P. steindachneri*) (Table 2). The comparison of anticodons for all tRNAs gene were resulted in Table S5. Most of the tRNAs were folded into classic clover-leaf secondary structures in *N. nigricans*, except for *trnS1* (GCT), which lacked the conventional dihydrouridine ‘DHU’ stem and loop (Fig. S2). The anticodon stem of *trnS1* (GCT) further formed a small loop instead of the conserved stem and loop structure. The G≡C and A=T bonding were conventionally observed in the secondary structures of tRNAs, however in six tRNAs; *trnA*(TGC), *trnN* (GTT), *trnY* (GTA), *trnS2* (TGA), *trnE* (TTC), and *trnP* (TGG) in *N. nigricans* shows G=T mismatches and forming weak bonds. Further, the *trnR* (TCG) and *trnH* (GTG) forms an unconventional small loop structure at 5′ end in the acceptor stem (Fig. S2). The total length of small and large rRNA genes of *N. nigricans* was 2593 bp and AT skewness was 0.307 and GC skewness was −0.180. In other Trionychidae species, the length of two rRNAs was varied from 2546 bp (*L. punctata*) to 2685 bp (*P. sinensis*). The AT skewness was varied from 0.268 (*P. sinensis*) to 0.329 (*T. triunguis*) and GC skewness was varied from −0.216 (*T. triunguis*) to −0.145 (*L. punctata*). The *rrnS* gene is located in between the *trnF* (GAA) and *trnV* (TAC); however, the *rrnL* gene is located in between *trnV* (TAC) and *trnL2* (TAA) in *N. nigricans* (Table 2).

### Control region (CR)

The length of CR in *N*. nigricans was 1290 bp and the A+T content was 68.52% (Table 2). Four tandem repeats; (ATTAT)_8_, (49bp)_2_, (AT)7, and (TATTA)20 with spacer of 151 bp between 1^st^ and 2^nd^ tandem repeat; 781 bp between 2^nd^ and 3^rd^ tandem repeat, and 87 bp between 3^rd^ and 4^th^ tandem repeat was observed (Fig. 4). The AT skew value was positive (0.07) and GC skew value was negative (−0.374). In the other Trionychidae species, the length of the CR varied from 1005 bp (*L. scutata*) to 1994 bp (*P. cantorii*). The AT and GC skew in other Trionychidae species have significant variation and ranges from −0.106 (*P. steindachneri*) to 0.067 (*T. triunguis*) and −0.453 (*P. cantorii*) to −0.230 (*P. steindachneri*) respectively. It is also resulted that the Adenine (A) composition is higher than Thymine (T) in *N. nigricans*, *C. indica*, *L. punctata*, *L. scutata*, *P. cantorii*, and *T. triunguis*. However, the Thymine (T) composition is higher than Adenine (A) is exceptionally resulted in *A. ferox*, *A. spinifera*, *D. subplana*, *N. formosa*, *P. steindachneri*, *P. sinensis*, and *R. swinhoei* (Table 2). The numbers of tandem repeats are higher at the 3′ end of the control region in most of the studied Trionychidae species. The number of tandem repeats in CRs varied from two (*A. spinifera*, *C. indica*, *R. swinhoei*, and *T. triunguis*) to eight (*P. sinensis*) and the AT composition was ranging from 61% (*P. sinensis*) to 70% (*N. formosa*). A single tandem repeat of AT was revealed in the two species (*L. scutata*, and *L. punctata*) under subfamily Cyclanorbinae. The length of the tandem repeats was also variable and ranges from 2 bp (*A. ferox*, *C. indica*, *D. subplana*, *L. scuata*, *L. punctata*) to 263 bp (*P. cantorii*). Exceptionally the overlapping tandem repeats were found in *A. ferox*, *P. cantorii*, and *P. sinensis*. The *A. ferox*, *A. spinifera*, *C. indica*, *R. swinhoei*, revealed a single tandem repeat (AT)_43_, (ATATTTAT)_13_, (AT)_27_ and (TATATATTA)_13_ respectively at the 3′ end of CR. Exceptionally, there is no tandem repeat present at the 5′ end of CR in *T. triunguis*, *L. scutata*, and *L. punctata*. Further, two tandem repeats, (TAAACACAC)_2_ and (ATCTATAT)_20_ with intergenic spacer of 71 bp was observed in *T. triunguis* and one tandem repeat each of (AT)_26_ and (AT)_28_ was observed in *L. scutata* and *L. punctata* respectively (Fig. 4). The repeated motifs within the CR, and duplication of CR within complete mitogenome; play an important role in regulating transcription and replication in the mitochondrial genome^36,37^. Thus, the resulted repeat sequence variations in *N. nigricans* and other Trionychidae species might be helpful to hypothesize the evolutionary pattern of this group.

**Figure 4.**
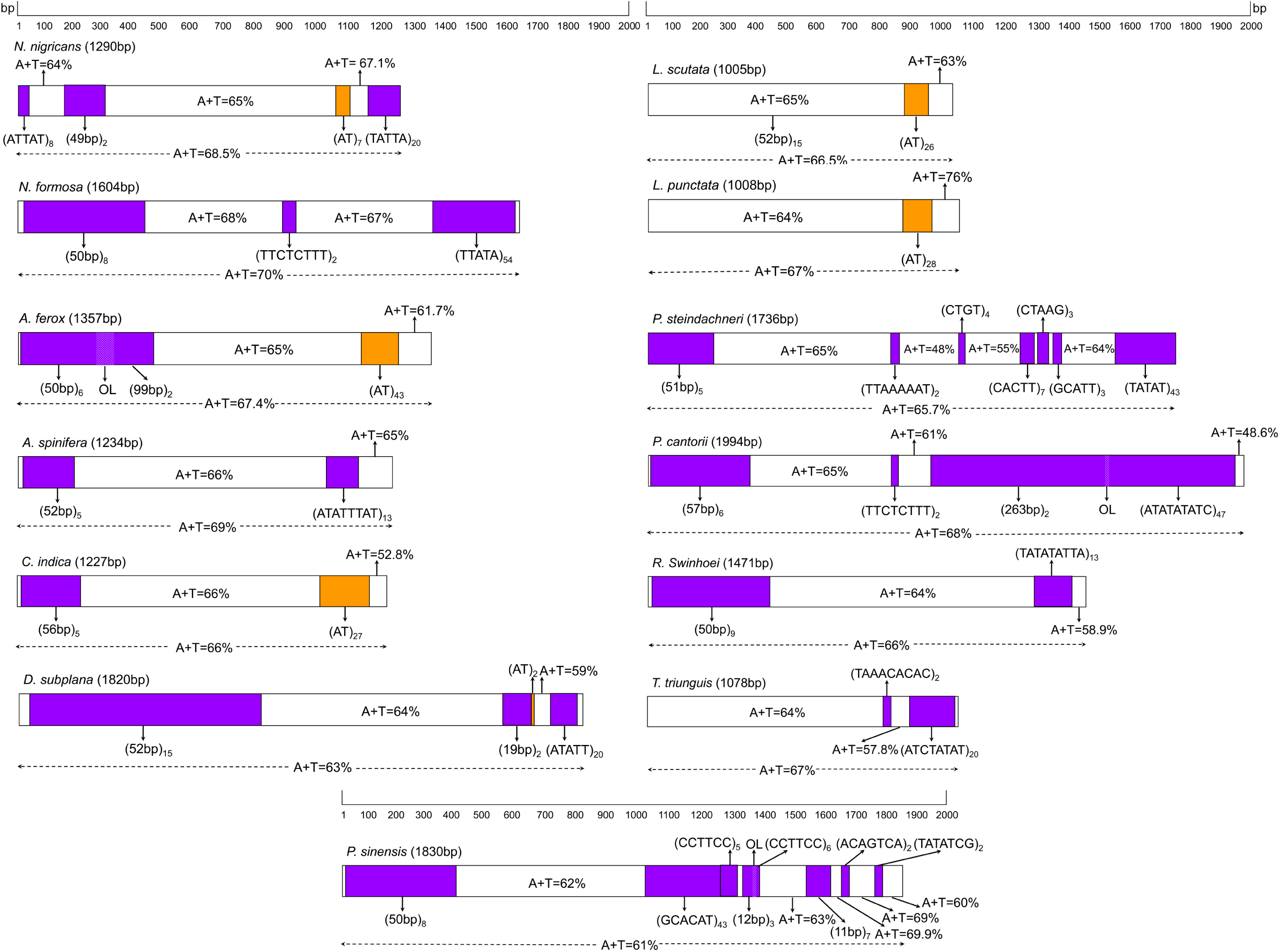
Comparison of control regions (CR) of the studied 13 Trionychidae species. Purple color boxes show the tandem repeats and orange color boxes show the AT tandem repeats within CR. OL= Overlapping regions. The length of the control regions were designated by representing scale. The tandem repeats in CR were predicted by the online Tandem Repeats Finder web tool (https://tandem.bu.edu/trf/trf.html) and edited manually in Adobe Photoshop CS 8.0.

### Synonymous and non-synonymous substitutions

The ratio of Ka/Ks was generally known as a pointer of selective pressure and evolutionary relations at the molecular level among homogenous or heterogeneous species^38,39,40^. It is reported that, the Ka/Ks>1 for positive selection, Ka/Ks=1 for neutral mutation, and Ka/Ks<1 for negative selection^41^. To investigate the evolutionary rates of *N. nigricans*, Ka/Ks substitutions were calculated and compared with other 10 Trionychidae species within 10 genera. The average Ka/Ks values of 13 PCGs were varied from 0.042 ± 0.013 (*cox3*) to 1.129 ± 0.274 (*nad4l*) and resulted the following order: *cox3*<*cox2*<*atp6*<*nad6*<*nad3*<*atp8*<*cox1*<*nad5*<*nad2*<*nad4*<*cytb*<*nad1*<*nad4l* (Table S6). Most of PCGs shows Ka/Ks values were <1, which indicated a strong negative selection among all Trionychidae species which reflects natural selection works against profitless mutations with negative selective coefficients as described earlier^40,42^. The average of synonymous and non-synonymous substitution variation were >1 in *cytb*, *nad1* and *nad4l* within all Trionychidae species including *N. nigricans*, with average value of Ka/Ks ratio 1.051, 1.120 and 1.129 respectively, which indicated the least selective pressure in *cytb*, *nad1* and *nad4l* genes. While the average of Ka/Ks were least in *cox3* (0.042), as compared with the lowest values of other species pairs, which indicated high selective pressure in *cox3* as compared with other PCGs (Fig. 5).

**Figure 5.**
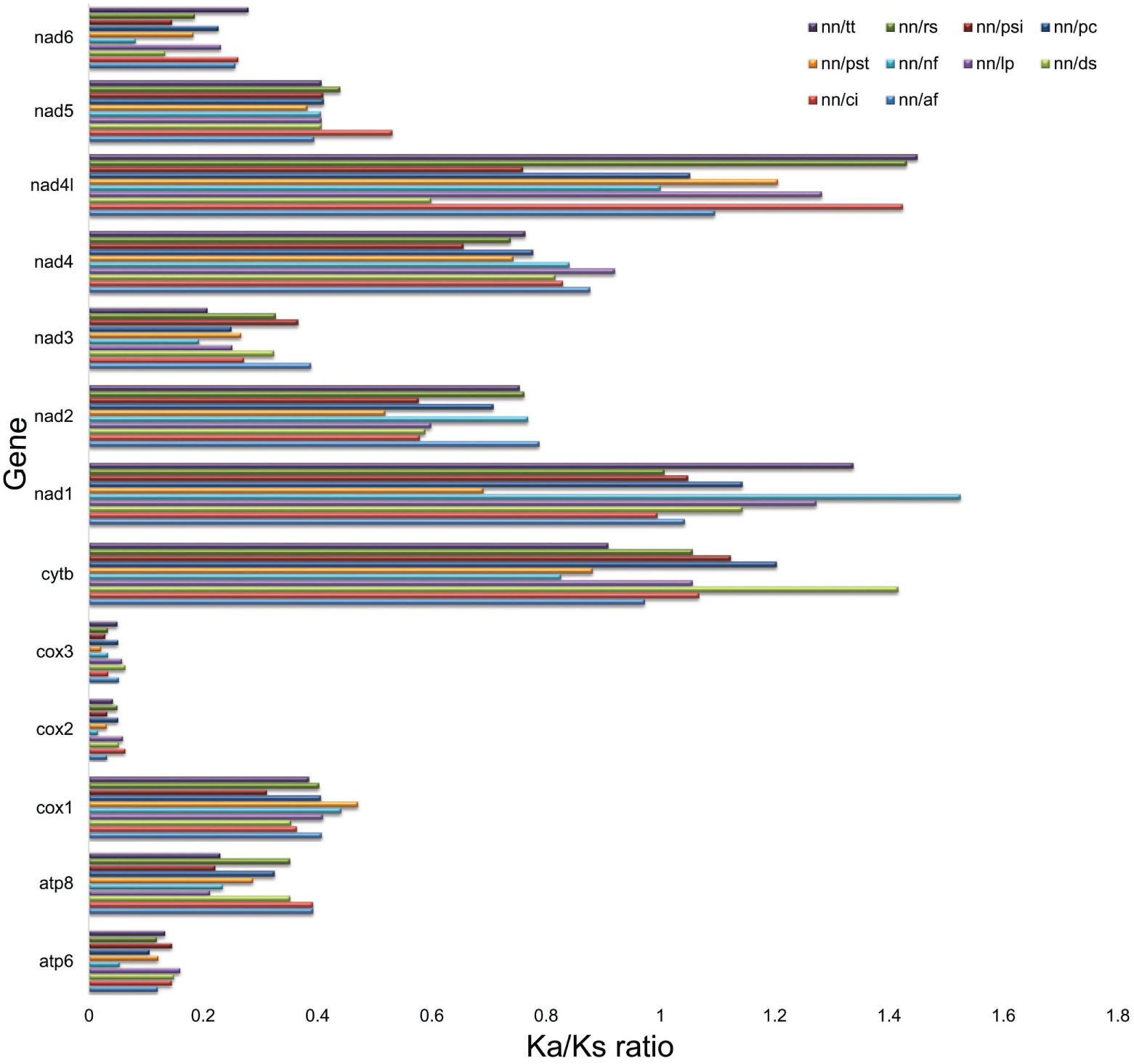
Ka/Ks ratios for the 13 mitochondrial PCGs among the references Trionychidae species. nn= *N. nigricans*, nf= *N. formosa*, af= *A. ferox*, ci= *C. indica*, ds= *D. subplana*, lp= *L. punctata*, pst= *P. steindachneri*, pc= *P. cantorii*, psi= *P. sinensis*, rs= *R. swinhoei*, and tt= *T. triunguis*. The software package DnaSPv5.0 was used to calculate the synonymous substitutions per synonymous sites (Ks) and non-synonymous substitutions per non-synonymous sites (Ka) and plotted manually in Microsoft Excel.

### Phylogenetic analysis

The aimed study constructed the phylogenetic relationships among the two suborders, seven families and 19 species of Testudines (Fig. 6). The phylogenetic tree revealed 13 species of Trionychidae family were clustered together with 100 bootstrap support and congruent with the previous phylogeny based hypothesis in Testudines systematics^1,2^. The Critically Endangered Black soft-shell turtle, *N. nigricans* resulted sister clade with *N. formosa*. The current phylogeny based on complete mitogenome, supports the previous classification of Trionychidae species. The 11 species shows distinct clade of three different tribes, Amydona, Apalonina, and Gigantaesuarochelys under Trionychinae subfamily. Further, the two species *L. punctata*, and *L. scutata* shows cohesive clustering as the members of subfamily Cyclanorbinae. The species of Geoemydidae and Testudinidae families were clustered together and shows sister clade of Platysternidae and Emididae species clade (Fig. 6). The Cheloniidae species (sea turtle) cladded separately in the ML tree. This phylogenetic analysis combined with all mitochondrial PCGs, is consistent with the previously described phylogeny using partial sequences of mitochondrial and nuclear gene^1,4,13,14,23^. Additionally, further taxon sampling from different taxonomic ranks and their mitogenomics data would be useful to reconcile the phylogenetic and evolutionary relationship of Testudines.

**Figure 6.**
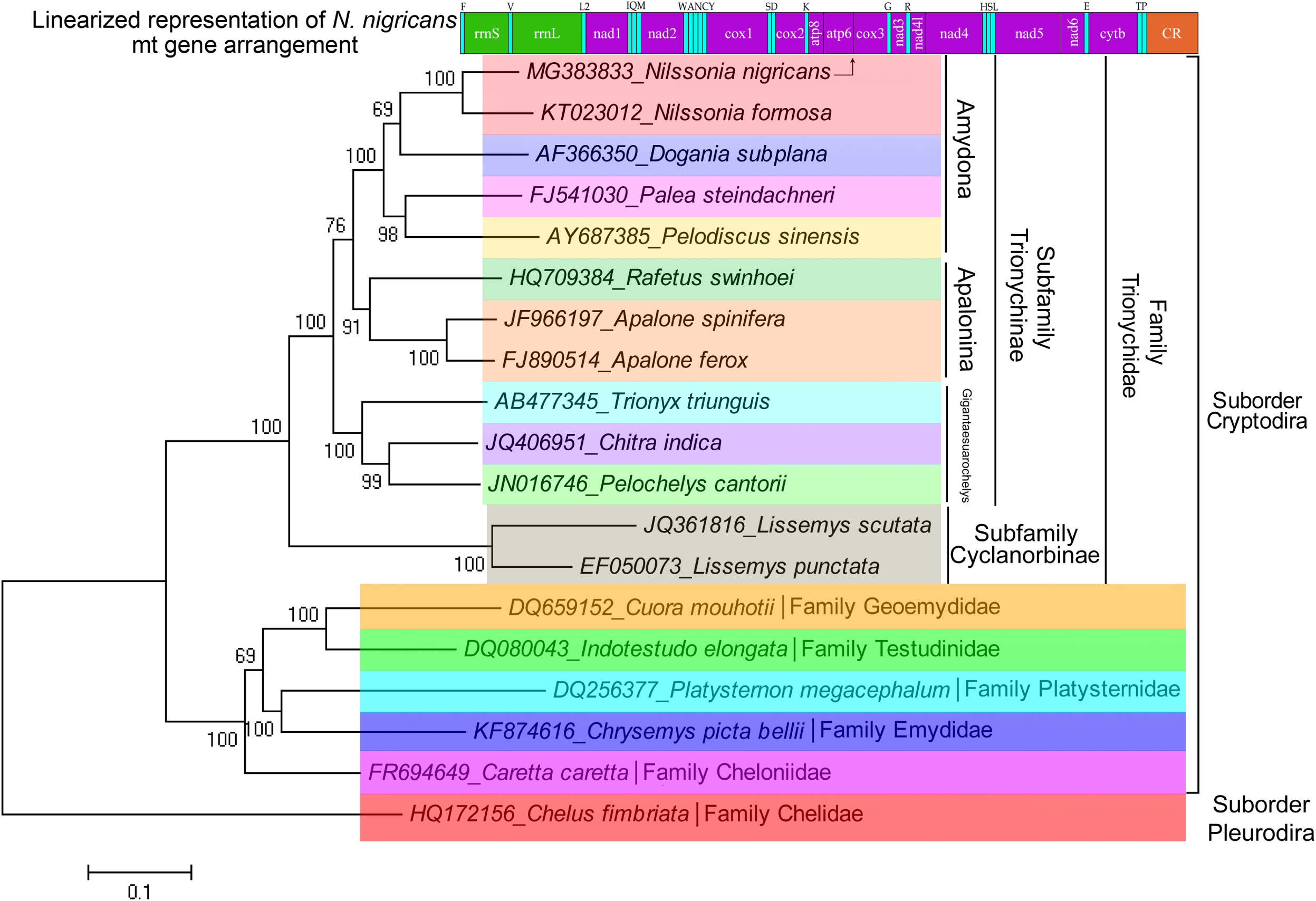
Maximum Likelihood phylogenetic tree based on the concatenated nucleotide sequences of 13 PCGs of the Trionychidae turtle species showing the evolutionary relationship of *N. nigricans*. Linearized represent of *N. nigricans* mitogenome gene arrangements also shown above the phylogenetic tree (blue and green color represents tRNA and rRNA genes respectively, purple and orange color represents PCGs and control region respectively). The established taxonomic ranks are incorporated with each clades by different colors. The figure was edited in Adobe Photoshop CS 8.0.

## Acknowledgements

We thank the Director of Zoological Survey of India (ZSI), Ministry of Environment, Forests and Climate Change (MoEF&CC), Govt. of India and Arunachal Pradesh Biodiversity Board for providing necessary permissions and facilities. We acknowledge the research funding for this work from DST-SERB National Post-Doctoral fellowship (F. No. PDF/2015/000302) to the first author (SK), ZSI, MoEF&CC in-house project, ‘National Faunal Genome Resources (NFGR)’ to the second author (VK). The funders had no role in study design, data collection and analysis or preparation of the manuscript.

## Author Contributions

S.K. and V.K. conceived and designed the experiment. S.K. collected specimens, performed taxonomic identification and captured photographs. V.K., K.T. and K.C. contributed chemicals. S.K., K.T., R.C. and D.S. generated DNA data. S.K., K.T. V.K., R.C., I.R. and A.P. analysed the data. S.K., I.R., K.T. and V.K. wrote the manuscript text and prepared the figures. All authors reviewed the manuscript.

## Additional Information

Supplementary information accompanies this paper at

## References

1. Engstrom, T.N., Shaffer, H.B. & McCord, W.P. Multiple Data Sets, High Homoplasy, and the Phylogeny of Softshell Turtles (Testudines: Trionychidae). Syst. Biol. 53, 693–710. (2004).

2. Meylan, P.A. The phylogenetic relationships of softshelled turtles (family Trionychidae). Bull. Nat. Hist. Mus. 186, 1–101. (1987).

3. Ernst, C.H., Altenburg, R.G.M. & Barbour, R.W. Turtles of the World. World Biodiversity Database. Version 1.2. Biodiversity Center of ETI, Amsterdam, CD-ROM. (2000).

4. Praschag, P., Hundsdörfer, A.K., Reza, A.H.M.A. & Fritz, U. Genetic evidence for wild-living *Aspideretes nigricans* and a molecular phylogeny of South Asian softshell turtles (Reptilia: Trionychidae: *Aspideretes*, *Nilssonia*). Zool. Scripta. 36, 301–310. (2007).

5. Anderson, J. Description of some new Asiatic Mammals and Chelonia. Ann. Mag. Nat. Hist. 16, 284–285. (1875).

6. van Dijk, P.P., Iverson, J.B., Shaffer, H.B., Bour, R. & Rhodin, A.G.J. Turtles of the world. update: annotated checklist of taxonomy, synonymy, distribution, and conservation status. Chelonian. Res. Monogr. 5, 000.165–000.242. (2011).

7. Rhodin, A.G.J., Iverson, J.B., Bour, R., Fritz, U., Georges, A., Shaffer, H.B. & van Dijk, P.P. Turtles of the World: Annotated Checklist and Atlas of Taxonomy, Synonymy, Distribution, and Conservation Status (8th Ed.). Turtle Taxonomy Working Group. 292 pp. ISBN: 978-1-5323-5026-9 (online). (2017).

8. Ahsan, M.F., Haque, M.N. & Fugler, C.M. Observations on Aspideretes nigricans, a semi-domesticated endemic turtle from eastern Bangladesh. Amphibia-Reptilia. 12, 131–136. (1991).

9. Khan, M.A.R. A ‘holy’ turtle of Bangladesh. Hornbill. 4, 7–11. (1980).

10. Rashid, S.M.A. The Aspideretes nigricans mystery. Brit. Herpetol. Soc. Bull. 34, 42–43. (1990).

11. IUCN. The IUCN red list of threatened species, Version 2017.3. (2018).

12. Le, M., Raxworthy, C.J., McCord, W.P. & Mertz, L. A molecular phylogeny of tortoises (Testudines: Testudinidae) based on mitochondrial and nuclear genes. Mol. Phylogenet. Evol. 40, 517–531. (2006).

13. Liebing, N., Praschag, P., Gosh, R., Vasudevan, K., Rashid, S.M.A., Rao, D.Q., Stuckas, H.U. & Fritz, U. Molecular phylogeny of the softshell turtle genus *Nilssonia* revisited, with first records of *N. formosa* for China and wild-living *N. nigricans* for Bangladesh. Vert. Zool. 62, 261–72. (2012).

14. Kundu, S., Laskar, B.A., Venkataraman, K., Banerjee, D. & Kumar, V. DNA barcoding of *Nilssonia* congeners corroborates existence of wild *N. nigricans* in northeast India. Mitochondrial DNA A DNA Mapp. Seq. Anal. 27, 2753–6. (2016).

15. Praschag, P. & Gemel, R. Identity of the black soft-shell turtle *Aspideretes nigricans* (Anderson, 1875), with remarks on related species. Faun. Abh. 23, 87–116. (2002).

16. Mindell, D.P., Sorenson, M.D., Dimcheff, D.E., Hasegawa, M., Ast, J.C. & Yuri, T. Interordinal Relationships of Birds and Other Reptiles Based on Whole Mitochondrial Genomes. Syst. Biol. 48, 138–152. (1999).

17. Parham, J.F., Feldman, C.R. & Boore, J.L. The complete mitochondrial genome of the enigmatic bigheaded turtle (Platysternon): description of unusual genomic features and the reconciliation of phylogenetic hypotheses based on mitochondrial and nuclear DNA. BMC Evol. Biol. 6, 1–11. (2006).

18. Zardoya, R. & Meyer, A. Complete mitochondrial genome suggests diapsid affinities of turtles. Proc. Natl. Acad. Sci. U.S.A. 95, 14226–14231. (1998).

19. Kumazawa, Y. & Nishida, M. Complete mitochondrial DNA sequences of the green turtle and blue-tailed mole skink: Statistical evidence for Archosaurian affinity of turtles. Mol. Biol. Evol. 16, 784–792. (1999).

20. Amer, S.A.M. & Kumazawa, Y. Complete sequence of the mitochondrial genome of the endangered nile soft-shelled turtle *Trionyx triunguis*. Egypt. J. Expt. Biol. (Zool). 5, 43–50. (2009).

21. Chen, X., Zhou, Z., Peng, X., Huang, X. & Chen, Z. Complete mitochondrial genome of the endangered Asian giant softshell turtle *Pelochelys cantorii* (Testudinata: Trionychidae). Mitochondrial DNA, 24, 111–113. (2013).

22. Jiang, J.J., Xia, E., Gao, C. & Gao, L. The complete mitochondrial genome of western painted turtle, *Chrysemys picta bellii* (Chrysemys, Emydidae). Mitochondrial DNA A DNA Mapp. Seq. Anal. 27, 787–788. (2016).

23. Li, H., Liu, J., Xiong, L., Zhang, H., Zhou, H., Yin, H., Jing, W. Li, J., Shi, Q., Wang, Y., Liu, J. & Nie, L. Phylogenetic relationships and divergence dates of softshell turtles (Testudines: Trionychidae) inferred from complete mitochondrial genomes. J. Evol. Biol. 30, 1011–1023. (2017).

24. Kearse, M., Moir, R., Wilson, A., Stones-Havas, S., Cheung, M., Sturrock, S., Buxton, S., Cooper, A., Markowitz, S., Duran, C., Thierer, T., Ashton, B., Mentjies, P., & Drummond, A. Geneious Basic: an integrated and extendable desktop software platform for the organization and analysis of sequence data. Bioinformatics, 28, 1647–1649. (2012).

25. Thompson, J.D., Gibson, T.J., Plewniak, F., Jeanmougin, F. & Higgins, D.G. The CLUSTAL_X windows interface: Flexible strategies for multiple sequence alignment aided by quality analysis tools. Nucl. Acids Res. 25, 4876–82. (1997).

26. Grant, J.R. & Stothard, P. The CGView Server: a comparative genomics tool for circular genomes. Nucl. Acids Res. 36, W181–W184. (2008).

27. Junqueira, A.C., Lessinger, A.C., Torres, T.T., Da, F.R., Vettore, A.L., Arruda, P. & Azeredo Espin A.M. The mitochondrial genome of the blowfly *Chrysomya chloropyga* (Diptera: Calliphoridae). Gene. 339, 7–15. (2004).

28. Tamura, K., Stecher, G., Peterson, D., Filipski, A. & Kumar, S. MEGA6: Molecular evolutionary genetics analysis Version 6.0. Mol. Biol. Evol. 30, 2725–9. (2013).

29. Lowe, T.M. & Chan, P.P. tRNAscan-SE On-line: Search and Contextual Analysis of Transfer RNA Genes. Nucl. Acids Res. 44, W54–57. (2016)

30. Laslett, D. & Canbäck B. ARWEN, a program to detect tRNA genes in metazoan mitochondrial nucleotide sequences. Bioinformatics. 24, 172–175. (2008).

31. Benson, G. Tandem repeats finder: A program to analyze DNA sequences. Nucl. Acids Res. 27, 573–580. (1999).

32. Librado, P., & Rozas, J. DnaSP v5: a software for comprehensive analysis of DNA polymorphism data. Bioinformatics. 25, 1451–1452. (2009).

33. Abascal, F., Zardoya, R. & Telford, M.J. TranslatorX: multiple alignment of nucleotide sequences guided by amino acid translations. Nucl. Acids Res. 38, W7–13. (2010).

34. Castresana J. Selection of conserved blocks from multiple alignments for their use in phylogenetic analysis. Mol. Biol. Evol. 17, 540–52. (2000).

35. Anderson, S., Bruijn, M., Coulson, A., Eperon, I., Sanger, F. & Young, I. Complete sequence of bovine mitochondrial DNA. Conserved features of the mammalian mitochondrial genome. J. Mol. Biol. 156, 683–717. (1982).

36. Hanna, Z.R., Henderson, J.B., Sellas, A.B., Fuchs, J., Bowie, R.C.K. & Dumbacher, J.P. Complete mitochondrial genome sequences of the northern spotted owl (*Strix occidentalis caurina*) and the barred owl (*Strix varia*; Aves: Strigiformes: Strigidae) confirm the presence of a duplicated control region. PeerJ. 5, e3901. DOI 10.7717/peerj.3901. (2017).

37. Bing, X., Fei, M., Yi, S. & Qing-Wei, L. Comparative Analysis of Complete Mitochondrial DNA Control Region of Four Species of Strigiformes. Acta. Genetica. Sin. 33, 965–974. (2006).

38. Shen, X., Ren, J., Cui, Z., Sha, Z., Wang, B., Xiang, J. & Liu, B. The complete mitochondrial genomes of two common shrimps (*Litopenaeus vannamei* and *Fenneropenaeus chinensis*) and their phylogenomic considerations. Gene. 403, 98–109. (2007).

39. Li, X.J., Huang, Y. & Lei, F.M. Comparative mitochondrial genomics and phylogenetic relationships of the Crossoptilon species (Phasianidae, Galliformes). BMC Genomics. 16, 42. (2015).

40. Zhu, K., Liang, Y., Wu, N., Guo, H., Zhang, N., Jiang, S. & Zhang D. Sequencing and characterization of the complete mitochondrial genome of Japanese Swellshark (*Cephalloscyllium umbratile*). Sci. Rep. 7, 15299. (2017).

41. Nei, M. & Kumar, S. Molecular Evolution and Phylogenetics. New York: Oxford University Press. (2000).

42. Yang, Z. & Bielawski, J. P. Statistical methods for detecting molecular adaptation. Trends Ecol. Evol. 15, 496–503. (2000).

